# Enhancing Hepatic MBOAT7 Expression Does Not Improve Nonalcoholic Steatohepatitis in Mice

**DOI:** 10.1101/2022.03.24.485677

**Authors:** Martin C. Sharpe, Kelly D. Pyles, Taylor Hallcox, Dakota R. Kamm, Michaela Piechowski, Bryan Fisk, Carolyn J. Albert, Danielle H. Carpenter, Barbara Ulmasov, David A. Ford, Brent A. Neuschwander-Tetri, Kyle S. McCommis

**Affiliations:** Biochemistry & Molecular Biology, Saint Louis University School of Medicine; Department of Internal Medicine, Division of Gastroenterology & Hepatology, Saint Louis University School of Medicine; Elizabeth H. and James S. McDonnell Genome Institute, Washington University School of Medicine; Pathology, Saint Louis University School of Medicine

**Keywords:** liver, NAFLD, NASH, MBOAT7, phosphatidylinositol, arachidonic acid, steatosis, fibrosis

## Abstract

**Background & Aims:** Genetic analyses of human NASH have revealed polymorphisms near the membrane bound O-acyl transferase domain containing 7 (*MBOAT7*) gene associated with worsened liver injury. NAFLD/NASH also appears to decrease MBOAT7 expression or activity independent of these polymorphisms. Thus, we hypothesized that enhancing MBOAT7 function in NASH would improve pathology.

**Approach & Results:** Male C57BL6/J mice were infected with adeno-associated virus 8 (AAV8) expressing MBOAT7 under control of the hepatocyte-specific thyroid hormone-binding globulin promoter, or control virus expressing green fluorescent protein (GFP). Mice were infected after NASH induction with either choline-deficient high-fat diet or Gubra Amylin NASH diet and compared to low-fat fed control mice. Both NASH diets increased liver weights, liver triglycerides, and plasma alanine and aspartate aminotransferase (ALT and AST) markers of liver injury, which were modestly yet significantly improved by MBOAT7 overexpression. However, NASH liver histology assessed by categorical scoring was not substantially improved by MBOAT7 overexpression. MBOAT7 regulates the formation of phosphatidylinositol (PI) predominantly by arachidonoylation of lysophosphatidylinositol (LPI). Shotgun lipidomics of NASH GFP-control livers suggested decreased MBOAT7 activity in that LPI content was elevated, and both total and arachidonoylated-PI were reduced. Surprisingly, MBOAT7 overexpression did not rescue the content of most arachidonoylated PI species but did normalize or increase the abundance of several oleate and linoleate-containing PI species. Free arachidonic acid was elevated but the MBOAT7 substrate arachidonoyl-CoA was found to be low in all NASH livers compared to low-fat fed mice, likely due to decreased expression of both long-chain acyl-CoA synthetases (ACSL) 1 and 4 in NASH livers compared to controls.

**Conclusions:** These results suggest MBOAT7 overexpression fails to measurably improve NASH pathology potentially due to insufficient abundance of its arachidonoyl-CoA substrate in fatty livers.

## Introduction

NAFLD and progression to NASH represent a large healthcare burden due the risk of progression to cirrhosis and lack of approved therapies (1, 2). In addition to being considered the hepatic manifestation of the metabolic syndrome, it is becoming more appreciated that genetic factors can play a significant role in NAFLD presence and progression (3, 4). One example is a common polymorphism (rs641738) in membrane bound O-acyltransferase domain containing 7 (*MBOAT7*) associated with increased NAFLD pathology (5–7). This *MBOAT7* rs641738 C>T variant is considered loss-of-function, resulting in decreased MBOAT7 expression (5).

MBOAT7 is a lysophosphatidylinositol (LPI) acyltransferase, combining an acyl group with LPI to form the diacyl phospholipid phosphatidylinositol (PI)(8). MBOAT7 appears to have selectivity for long-chain polyunsaturated fatty acids such as arachidonic acid (8, 9), meaning that while total PI levels may not be altered, the abundance of specific unsaturated PI species are significantly regulated by MBOAT7.

In agreement with human genetic studies, cell culture and mouse models suggest that MBOAT7 deficiency leads to hepatocellular lipid accumulation and increased liver injury and fibrosis (10–14). Several reports suggest this increase in hepatic triglyceride with loss of MBOAT7 stems from enhanced sterol regulatory element binding protein 1 (SREBP1) expression/activity and increased *de novo* lipogenesis (11, 14), while another study suggests a route for PI conversion to diacylglycerol then triglyceride (12).

It was reported that obesity, presumably associated with NAFLD, decreases hepatic *MBOAT7* expression independent of the rs641738 polymorphism (10). Based on these previous human, rodent, and cell culture studies of MBOAT7 deficiency, we hypothesized that enhancing MBOAT7 expression/activity would improve NASH pathology. To test this, we stably overexpressed MBOAT7 by adeno-associated viral infection in two different dietary models of murine NASH.

## Experimental Procedures

### Human Gene Expression and Metabolomic Database Analysis

We mined publicly available Gene Expression Omnibus (GEO) Datasets for studies of human liver microarray and RNA sequencing expression profiles in NAFLD and/or NASH and extracted expression data for *MBOAT7*. Data for *MBOAT7* expression was combined from GSE89632 (15), GSE126848 (16), GSE135251 (17), GSE167523 (18), and GSE163211 (19), and normalized to healthy control liver expression levels. In total, *MBAOT7* expression was measured in livers from individuals with biopsy-determined status as normal control (NC, n=38), healthy obese (HO, n=98), “simple steatosis” or NAFL (n=225), or NASH with varying degrees of fibrosis, including NASH cirrhosis (n=391). We also mined publicly available lipidomic analyses for PI measured in human NAFLD/NASH compared to control livers. These included data deposited in Metabolomics Workbench ST000915 (20), as well as supplemental data from an open access publication (21). Total PI levels as well as the predominant arachidonoylated PI species (38:4), were assessed in a total of 80 NC livers, 51 HO livers, 160 NAFL livers, and 134 NASH livers. Since the two studies were performed using different mass spectrometry methods and normalized in a different manner (protein concentration vs starting tissue weight), data for each study was normalized to NC prior to combining.

### Animals

Animal care and experimentation was approved by the IACUC of Saint Louis University and complied with criteria outlined in the *Guide for the Care and Use of Laboratory Animals*. Five-week-old male C57BL/6J mice were purchased from Jackson Laboratories (000664, The Jackson Laboratories, Bar Harbor, ME). Mice were housed in specific-pathogen-free, climate-controlled rooms with a 6:00-18:00 light ON/OFF cycle. Mice were group housed, up to 5 per cage, with *ad libitum* access to water and food. At 6-weeks of age, mice were randomly chosen to consume either low fat (LF) diet (D12450K, Research Diets, New Brunswick, NJ; 10% kcal fat, 20%kcal protein, and 70%kcal carbohydrate) or choline-deficient aminoacid defined high-fat diet (CDAHFD)(Research Diets A06071309; 46%kcal fat, 18%kcal protein, and 36%kcal carbohydrate). After consuming diets for 6 weeks, CDAHFD-fed mice were infected with 7×10^11^ genome copies of adeno-associated virus serotype 8 (AAV8) via intravenous injection of the tail vein. AAV8 was purchased from Vector Biolabs (Malvern, PA), and mice were randomly chosen to be infected with AAV8 expressing either a control green fluorescent protein (GFP; catalog VB1743) or murine *Mboat7* (RefSeq BC023417, catalog AAV-264332), both under the control of the hepatocyte-specific thyroxine binding globulin promoter. Mice continued to consume special diets for another 4 weeks at which point they were fasted for 2-3 hours, and euthanized by CO2 asphyxiation. Blood was collected from the abdominal aorta into an EDTA coated tube and spun at 2,000 x g for 10 minutes to collect plasma. Liver was excised, weighed, divided, and either snap frozen in liquid nitrogen, fixed in 10% neutral buffered formalin, or placed in 1 mL of RNALater (Ambion, Austin, TX). A second set of 6-week old mice was fed either LF diet (D12450K) or Gubra Amylin NASH diet (GAN)(Research Diets D09100310; 40%kcal fat from mostly palm oil, 20%kcal protein, and 40%kcal carbohydrate containing fructose, as well as 2%wt cholesterol). After 19-weeks on diet, mice were randomized to infection with AAV8-GFP or AAV8-MBOAT7 viral infection as described above. 6 weeks after infection, mice were fasted for 4 hours and subjected to an i.p. glucose tolerance test (GTT) (1mg/kg body weight glucose), and blood glucose was measured by lateral nick of the tail vein and collecting a drop of blood with a Contour Next EZ glucometer (Ascensia Diabetes Care, Parsippany, NJ). These mice consumed diets for 28 weeks total, then euthanized by CO_2_ asphyxiation after a 2-3 hour fast and plasma/tissue processed as described above.

### Gene Expression Analyses

Total RNA was isolated from ~5 mg frozen liver tissue by homogenization in 1 mL RNA-STAT (TelTest, Friendswood, TX) with isopropanol and EtOH precipitation and analyzed as performed previously (22). Target gene Ct values were normalized to reference gene (*Rplp0*) Ct values by the 2^-ΔΔCt^ method. Oligonucleotide primer sequences are listed in Supplemental Table 1.

### Protein Expression Analyses

Protein extracts were prepared by homogenizing ~50 mg liver tissue in lysis buffer 15 mM NaCl, 25 mM Tris base, 1 mM EDTA, 0.2% NP-40, and 10% glycerol), supplemented with protease and phosphatase inhibitors. To assess MBOAT7 localization, liver samples were fractionated by dounce homogenization and centrifuged 1,000g x 5 min @ 4°C to pellet whole cells and nuclei, and then the supernatant centrifuged at 21,100g x 20 min @ 4°C to separate membrane and cytosolic fractions. Protein concentration was measured by a MicroBCA kit (ThermoFisher Scientific, Waltham, MA). 50 μg of protein was electrophoresed on Criterion 4-15% precast polyacrylamide gels (Bio-Rad, Hercules, CA) and transferred onto polyvinyldifluoride membranes. Protein lysates were not boiled prior to electrophoresis for MBOAT7 immunoblots. Membranes were blocked for at least 1 hour in 5% bovine serum albumin (BSA)(MilliporeSigma, Burlington, MA) in tris-buffered saline with Tween-20 (TBST). Membranes were incubated with primary antibodies at 1:1000 dilution overnight at 4°C. Membranes were then washed 5 x 5 min in TBST, and then incubated with secondary antibodies at 1:10,000 dilution in 5%-BSA-TBST for 1 hour. After washing 5 x 5 min in TBST, membranes were developed on an Odyssey imaging system and analyzed with Image Studio software (Li-Cor Biosciences, Lincoln, NE). Information for all antibodies is listed in Supplemental Table 2.

### Plasma Insulin, Lipids, and Hepatic Triglyceride Analyses

Plasma insulin, cholesterol, triglyceride (TAG), and non-esterified fatty acids (NEFA) were measured with commercial assays as previously described (23). Liver TAG concentrations were measured as previously described (23) by homogenizing ~100 mg frozen liver in saline to provide 0.1 mg/μL. This liver homogenate was combined 1:1 with 1% sodium deoxycholate, vortexed, and incubated at 37°C for 5 min to solubilize lipids. TAG was then measured by colorimetric assay (TR22421, ThermoFisher Scientific, Waltham, MA).

### Liver Histology and Scoring

Formalin-fixed liver was embedded in paraffin blocks and sectioned onto glass slides. Slides were stained by H&E and picrosirius red and were evaluated by a histopathologist blinded to diet and treatment groups. Slides were scored for steatosis, inflammation, and fibrosis using typical NAFLD activity score (NAS) criteria (24). In addition, total percentage of hepatocytes with steatosis, and stratification of macrosteatosis or microsteatosis was recorded. Percentage of picrosirius red-stained fibrosis was also quantified digitally by averaging the red-stained area from at least 10 independent pictures of each slide using Image J software as performed previously (22).

### Liver Lipidomic Analyses

Lipids were extracted by pulverizing ~100 mg frozen liver and subjecting ~15 mg tissue to a modified Bligh-Dyer extraction as performed previously (25) after spiking in a cocktail of internal standards including di-20:0 phosphatidylcholine (PC), di-14:0 phosphatidylethanolamine (PE), di-14:0 phosphatidylserine (PS), N17:0 sphingomyelin, 17:0 cholesteryl ester, 17:0 fatty acid, N17:0 ceramide, 17:0 lysophosphatidylcholine (LPC), 17:0-18:1-d_5_ phosphatidylinositol (PI; 850111, Avanti Polar Lipids, Alabaster, AL), and 17:0-d_5_ lysophosphatidylinositol (LPI; 850108, Avanti Polar Lipids, Alabaster, AL). Lipid extracts were diluted in methanol/chloroform (4/1, v/v) and lipid species quantified using shotgun lipidomics by electrospray ionization-mass spectrometry on a triple-quadrupole Quantum Ultra (ThermoFisher Scientific, Waltham, MA) as performed previously (26). Individual molecular species were quantified by comparing the ion intensities of the individual molecular species to that of the lipid class internal standard with additional corrections for type I and type II ^13^C isotope effects. 38:4 and 36:4 PI were confirmed as 18:0/20:4 and 16:0/20:4 by product ion scanning for individual fatty acid constituents for oleate, palmitate, and arachidonate at *m/z* of 281.3, 255.2, and 303.4, respectively, at collision energy of +35 eV.

Measurement of free fatty acids was performed as previously described (25). Briefly, 50 μL lipid extract was dried and resuspended in 100 μL 2.5% diisopropylethylamine and 5% pentafluorobenzyl bromide in acetonitrile. Samples were incubated at 45°C for 1 hour, allowed to cool at room temp., and twice dried and resuspended in 1 mL ethyl acetate. After drying one final time samples were suspended in 200 μL ethyl acetate and free fatty acids detected by gas chromatography/mass spectrometry and selected ion monitoring as previously performed (25).

Acyl-CoAs were measured by homogenizing ~100 mg frozen liver in water (0.25 g/mL), and protein precipitation performed to extract acyl-CoAs from 50 μL of homogenate in the presence of a d4-palmitoyl-CoA internal standard. Analysis of acyl-CoA was performed with a high-performance liquid chromatography system (Shimadzu 20A, Kyoto, Japan) coupled to a 6500QTRAP+ mass spectrometer (AB Sciex LLC, Framingham, MA) operated in positive multiple reaction monitoring mode. Data processing was conducted with Analyst 1.6.3 software. Quality control (QC) samples were prepared by pooling aliquots of study samples and this QC sample was injected between every ten study samples to monitor instrument performance. Only acyl-CoA species with coefficient of variance <15% of QC injections are reported. Relative quantification of acyl-CoA is reported as the peak area ratios of the analytes to the corresponding internal standard.

### Statistical Analyses

Unless stated, data are presented as individual data points with mean ± S.E.M. All datasets were analyzed for statistical significance by one-way ANOVA with post-hoc analysis by Tukey’s correction for multiple comparisons using GraphPad Prism Version 9.3.1 or RStudio. Adjusted *p* values <0.05 were considered significant.

## Results

### Human NAFLD and NASH are Associated with Decreased MBOAT7 Expression and Activity

A previous report observed dramatically decreased hepatic *MBOAT7* expression in a small set of obese individuals, presumably with NAFLD, compared to lean controls (10). To assess if this decrease in *MBOAT7* held true in larger cohorts of patients with confirmed NAFLD or NASH, we mined GEO datasets for *MBOAT7* expression. Indeed, livers from obese patients without steatosis (healthy obese, HO), or patients with biopsy confirmed NAFL or NASH all displayed significantly reduced MBOAT7 compared to livers from lean normal controls (NC) (Fig. 1A). MBOAT7 transfers an acyl group, thought to be predominantly arachidonic acid (8, 9), to lysophosphatidylinositol (LPI) to form phosphatidylinositol (PI) (Fig. 1B). To investigate MBOAT7 activity in human NAFLD we also mined available lipidomic studies in which hepatic PI was measured. The predominant PI species in the liver contains arachidonic acid (20:4), and this 38:4 PI was significantly decreased in HO, NAFL, and NASH livers compared to NC (Fig. 1C), suggesting decreased MBOAT7 activity. Interestingly, both 38:4 PI and Total PI levels displayed a significant negative association with the total NAFLD activity score (NAS) (Fig. 1D and E), suggesting that decreased hepatic MBOAT7 activity and PI levels associate with worsened NAFLD pathology. Altogether, these results suggest that obesity and NAFLD are associated with reduced hepatic MBOAT7 gene expression and activity.

**FIGURE 1.**
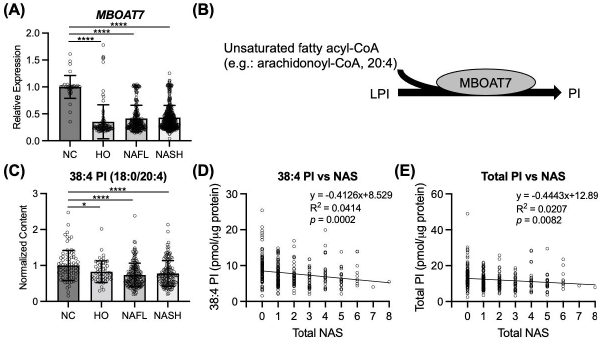
Human NAFLD/NASH decreases hepatic *MBOAT7* expression and activity. (A) Hepatic *MBOAT7* expression combined from GSE89632, GSE126848, GSE135251, GSE167523, and GSE163211 reveals decreased *MBOAT7* in humans with biopsy confirmation of healthy obesity (no steatosis; HO, *n*=98), “simple steatosis” or NAFL (*n*=225), or NASH (*n*=391) compared to lean healthy control (HC, *n*=38) livers. (B) Schematic showing the catalytic activity of MBOAT7 combining lysophosphatidylinositol (LPI) and a fatty acyl-CoA to form phosphatidylinositol (PI). (C) The abundant, arachidonoylated PI species (18:0/20:4), combined from (20, 21), is decreased in HO, NAFL, and NASH compared to NC livers (*n*=51, 160, 134, and 80, respectively). (D) The MBOAT7 product, 38:4 PI, measured in (21), is negatively associated with total NAFLD activity score (NAS). (E) Total PI, measured in (21), is negatively associated with total NAS. Data presented as mean ± SD, analyzed by one-way ANOVA with Tukey’s post-hoc correction for multiple comparisons, \**p* < 0.05, \*\*\*\**p* < 0.0001.

### Hepatic MBOAT7 Overexpression in Mice Modestly Improves Plasma and Gene Expression Markers for NASH Liver Injury

If reduced MBOAT7 expression or activity contributes to NAFLD pathology, we hypothesized that increasing MBOAT7 expression may improve NASH. To test this, we used two different dietary models of murine NASH, and infected with adeno-associated virus to stably express MBOAT7 or GFP control in the liver. AAV8-TBG-MBOAT7 significantly enhanced hepatic *Mboat7* RNA (Fig. 2A and J) and MBOAT7 protein expression (Fig. 2B,C,K,L) compared to both low-fat (LF)-fed control mice and mice fed NASH diets treated with AAV8-TBG-GFP control virus.

**FIGURE 2.**
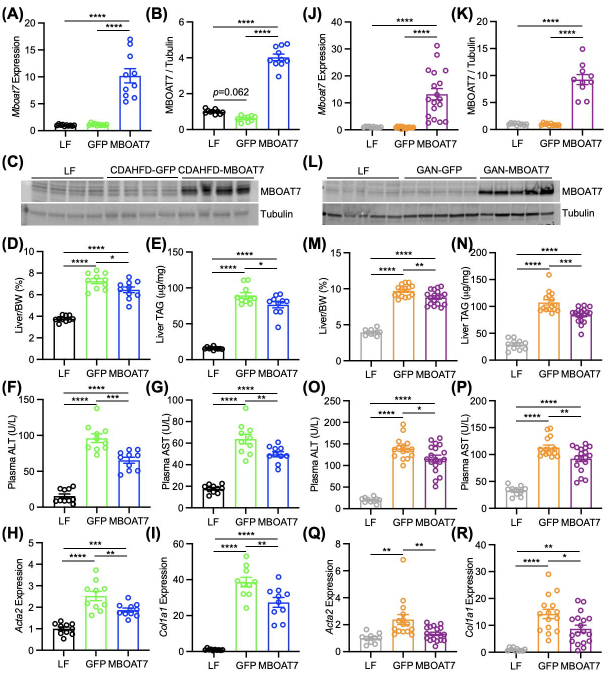
MBOAT7 overexpression in mice modestly improves hepatic triglycerides, and plasma and gene expression markers of NASH liver injury. (A) Mice were fed either low fat (LF) control diet or choline-deficient high-fat diet (CDAHFD) (A-I) or Gubra Amylin NASH diet (GAN) (J-R) to induce NASH. Mice consuming NASH diets were infected with adeno-associated viruses to express either control green fluorescent protein (GFP) or MBOAT7. (A, J) Hepatic *Mboat7* gene expression overexpressed by AAV8-MBOAT7 in CDAHFD and GAN models, respectively. (B-C, K-L) Hepatic MBOAT7 expression in CDAHFD and GAN models, respectively (*n*=10). (D, M) Liver weights normalized to body weight (BW) is modestly but significantly decreased by MBOAT7 overexpression. (E, N) Liver triglyceride (TAG) concentrations modestly yet significantly decreased by MBOAT7 overexpression. (F-G, O-P) Plasma levels of the liver injury markers alanine transaminase (ALT) and aspartate transaminase (AST) were significantly improved by MBOAT7 overexpression. (H-I, Q-R) Hepatic gene expression of *Acta2* and *Col1a1* significantly reduced in mice with MBOAT7 overexpression. Data presented as mean ± SEM, CDAHFD study: *n*=10 each; GAN study: LF n=10, GFP n=15, MBOAT7 n=18. Data analyzed by one-way ANOVA with Tukey’s post-hoc correction for multiple comparisons, \**p* < 0.05, \*\**p* < 0.001, \*\*\**p* < 0.001, \*\*\*\**p* < 0.0001.

As expected, CDAHFD did not induce obesity compared to LF diet (Fig. S1A), and actually decreased glycemia compared to LF-fed mice (Fig. S1B). Conversely, GAN diet induced obesity, slight hyperglycemia, a trend for increased insulinemia, as well as significantly reduced glucose tolerance compared to LF-fed mice (Fig. S1C-G). GAN diet significantly elevated plasma cholesterol levels, but significantly reduced plasma TAG and NEFA compared to LF diet, and MBOAT7 had no effect on these plasma lipids compared to GFP control (Fig. S1H-J). Thus, while GAN diet produced modest obesity and metabolic dysregulation, hepatic MBOAT7 overexpression did not alter these measures.

Both NASH diets produced significant hepatomegaly (Fig. 2D and M) and hepatic triglyceride accumulation (Fig. 2E and N), which were both modestly but significantly improved by MBOAT7 overexpression in both diet models. Likewise, the plasma markers for liver injury, ALT and AST, were elevated by both NASH diets and significantly improved by MBOAT7 overexpression (Fig. 2F,G,O,P). Lastly, hepatic gene expression markers of stellate cell activation and fibrogenesis were significantly elevated by both models of NASH, yet while *Acta2* and *Col1a1* were significantly reduced (Fig. 2H,I,Q,R), *Col3a1* and *Timp1* were not improved by MBOAT7 overexpression (Fig. S1K,L,M,N). Altogether, these results suggest that hepatic MBOAT7 overexpression may have beneficial effects on NASH pathology.

### Hepatic MBOAT7 Overexpression Does Not Measurably Improve Liver Histology

Histologic analyses confirmed the presence of extensive steatosis, inflammation, and fibrosis in both models of NASH compared to livers from LF-fed mice (Fig. 3A and B). Histologic scoring revealed that MBOAT7 overexpression did not improve any of these NAFLD histology indices in either NASH diet model (Fig. 3C and F). Additionally, the percentage of hepatic steatosis was estimated, and characterized as either macrovesicular or microvesicular lipid droplets. The CDAHFD NASH model produced almost entirely macrovesicular steatosis (Fig. 3A and D), however the GAN diet resulted in nearly 40% microvesicular steatosis which was significantly reduced by MBOAT7 overexpression (Fig. 3B and G). Digital quantification of the picrosirius red staining validated the histology scores in that both NASH diets significantly induced fibrosis, which was not improved by MBOAT7 overexpression (Fig. 3E and H). In summary, hepatic MBOAT7 overexpression did not markedly improve NASH pathology.

**FIGURE 3.**
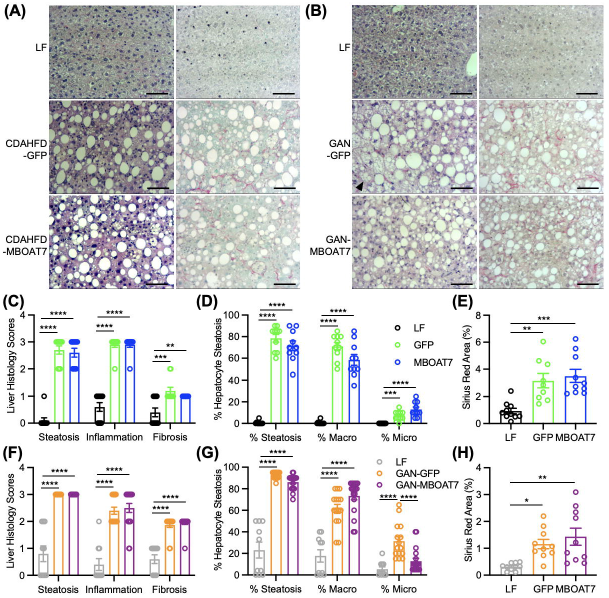
MBOAT7 overexpression does not substantially improve NASH histology. (A,B) Representative H&E and picrosirius red stains of livers from mice fed LF or CDAHFD or GAN diet, respectively (10X magnification, scale bar = 100 μm). (C,F) Liver histology using broad categorical scoring for steatosis, inflammation, and fibrosis in CDAHFD and GAN models, respectively suggests no improvement in histology with MBOAT7 overexpression. (D,G) Histological assessment of % of hepatocytes with any steatosis, or macro-vs microsteatosis indicates that GAN diet induces a notable amount of microsteatosis which is significantly reduced by MBOAT7 overexpression. (E,H) Digitally quantified sirius red stained area suggests no improvement in fibrosis with MBOAT7 overexpression (all *n*=10). Data presented as mean ± SEM, CDAHFD study: *n*=10 each; GAN study: LF n=10, GFP n=15, MBOAT7 n=18. Data analyzed by one-way ANOVA with Tukey’s post-hoc correction for multiple comparisons, \**p* < 0.05, \*\**p* < 0.001, \*\*\**p* < 0.001, \*\*\*\**p* < 0.0001.

### MBOAT7 Overexpression Alters Various Hepatic Phosphatidylinositol Levels

To determine if MBOAT7 overexpression enhanced MBOAT7 activity, we measured liver LPI and PI concentrations by shotgun lipidomics. If NASH decreases MBOAT7 activity, one would expect an accumulation of LPI, and indeed several LPI species (Fig. 4A and D) as well as total LPI levels (Fig. S2A and D) were increased by both NASH diets compared to LF. MBOAT7 overexpression would be predicted to reduce this accumulation of LPI substrate, yet to our surprise, total LPI levels were not affected by MBOAT7 (Fig. S2A and D), which was driven by the abundant 18:0 LPI not being reduced by MBOAT7 overexpression (Fig. 4A and D). However, a few less abundant LPI species, including 16:0 and 20:2 in the CDAHFD model and 16:0 in the GAN model were significantly reduced by MBOAT7 overexpression (Fig. 4A and D). Conversely, reduced MBOAT7 activity in NASH is expected to reduce PI levels, and indeed the majority of individual PI species (Fig. 4B,C,E,F) and total PI (Fig. S2B and E) were significantly reduced by both NASH diets compared to LF. Arachidonate-containing 38:4 PI accounts for roughly half of total PI in the liver. Since MBOAT7 is suggested to be a relatively specific acyltransferase for arachidonic acid (8, 9), we expected this species to be normalized by MBOAT7 overexpression. Surprisingly, 38:4 was not rescued by MBOAT7 in either NASH model and was even significantly decreased by MBOAT7 overexpression in the GAN diet (Fig. 4B and E). Another more abundant arachidonate-containing PI specie, 36:4, was also not increased with MBOAT7 overexpression in either NASH diet (Fig. 4C and F). However, many lesser-abundant PI species, including non-arachidonate containing species, were significantly elevated by MBOAT7 overexpression in both NASH models (Fig. 4C and F). Indeed, if the predominant 38:4 PI is excluded, total PI levels were significantly increased by MBOAT7 overexpression (Fig. S2C and F). These data suggest that while MBOAT7 did not rescue the major arachidonate-containing PI species, other PI levels were significantly elevated by MBOAT7 overexpression in both NASH models.

**FIGURE 4.**
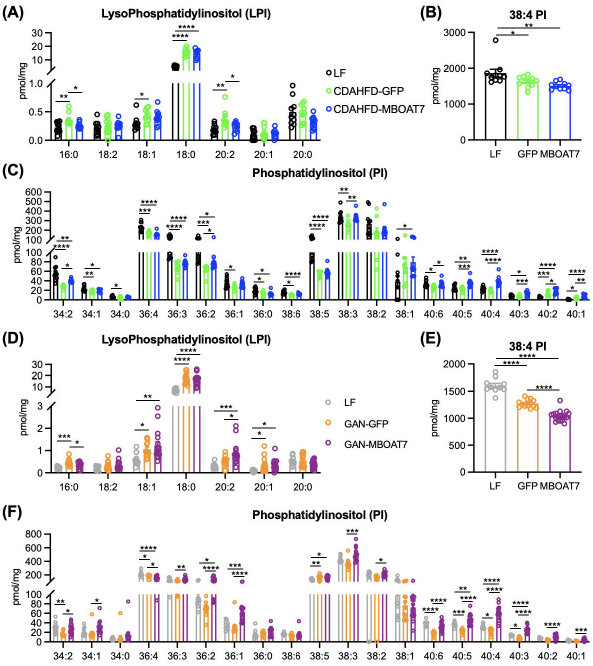
MBOAT7 overexpression increases many phosphatidylinositol species. (A,D) Hepatic lysophosphatidylinositol (LPI) measured in CDAHFD and GAN models, respectively, suggest that NASH induces increased abundance of LPI which is largely unaltered by MBOAT7 overexpression. (B,E) The most abundant phosphatidylinositol (PI) species (38:4) is reduced in the CDAHFD and GAN models of NASH, respectively, and not rescued by MBOAT7 overexpression. (C,F) PI species are almost all significantly reduced the CDAHFD and GAN models of NASH, respectively, and many are significantly increased by MBOAT7 overexpression in both models. Data presented as mean ± SEM, CDAHFD study: *n*=10 each; GAN study: LF n=10, GFP n=15, MBOAT7 n=18. Data analyzed by one-way ANOVA with Tukey’s post-hoc correction for multiple comparisons, \**p* < 0.05, \*\**p* < 0.001, \*\*\**p* < 0.001, \*\*\*\**p* < 0.0001.

Due to the lack of effect on the predominant arachidonate-containing PI species, we next tested whether the overexpressed MBOAT7 localized properly to endomembranes (endoplasmic reticulum, mitochondrial-associated membranes, and lipid droplets)(5, 27). In both NASH models, the majority of overexpressed MBOAT7 was localized to the membrane fraction containing the endoplasmic reticulum marker calnexin (Fig. S3A and B). Only small amounts of the overexpressed MBOAT7 appeared in the cytosolic fraction marked by lactate dehydrogenase (Fig. S3A and B). Thus, the overexpressed MBOAT7 was properly localized to endomembranes, allowing it to increase the abundance of many PI species but not the major arachidonate-containing PIs.

Our shotgun lipidomic analyses also measured hepatic lysophosphatidylcholine, phosphatidylcholine, phosphatidylethanolamine, phosphatidylserine, cholesteryl esters, ceramides, and sphingomyelins. Both NASH diets resulted in many significant changes in these lipids compared to LF control livers (Fig. S4-7). As MBOAT7 has been previously described to have specific LPI acyltransferase activity, unsurprisingly there were extremely few significant changes between GFP control and MBOAT7 overexpression. Specifically, only 36:0 phosphatidylcholine, 16:1 and 16:0 cholesteryl esters, and 16:1 sphingomyelin were slightly but significantly altered by MBOAT7 in the GAN diet model (Fig. S6 and 7). Altogether, these results suggest that as expected, MBOAT7 overexpression predominantly affected the abundance of PI.

### Murine NASH Decreases Arachidonic Acid Availability via Decreased Long-chain Acyl-CoA Synthetase Expression

While reviewing phospholipid abundances, we noted that the primary arachidonate-containing 38:4 and 36:4 species of both phosphatidylethanolamine and phosphatidylserine were significantly reduced by both NASH diets compared to LF diet livers (Fig. S4C and D and S6C and D). We hypothesized that arachidonic acid levels may be limited in NASH, helping to explain why MBOAT7 overexpression did not increase the main arachidonate-containing PI species. We first measured hepatic free fatty acid levels which uncovered an increase in 20:4 arachidonic acid in both NASH diets compared to LF diet (Fig. 5A and E). MBOAT7 overexpression did not affect this increase in arachidonic acid (Fig. 5A and E), and overall, total free fatty acid levels were not greatly altered in NASH livers compared to LF or by MBOAT7 compared to GFP overexpression (Fig. 5B and F). However, for arachidonate to become a substrate of MBOAT7, it needs to be activated by condensation with CoA, converting it into arachidonoyl-CoA. Therefore, we also measured long-chain acyl-CoAs in these livers, and 20:4 arachidonoyl-CoA, as well as most other acyl-CoAs, was significantly decreased by NASH diets compared to LF (Fig. 5C,D,G,H). MBOAT7 overexpression had no effect on arachidonoyl-CoA level compared to GFP control livers, therefore all NASH livers displayed relatively limited amounts of arachidonoyl-CoA, potentially explaining why MBOAT7 failed to enhance the abundance of the arachidonoylated 38:4 and 36:4 PI species.

**FIGURE 5.**
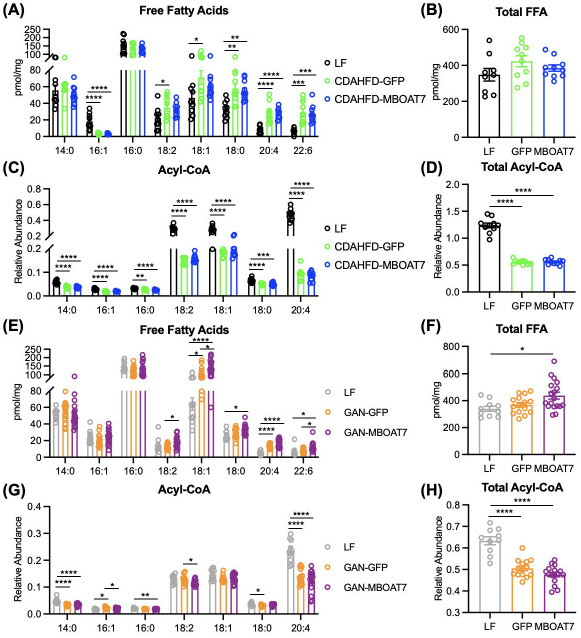
NASH decreases hepatic arachidonoyl-CoA concentrations. (A,E) Hepatic free fatty acid abundance in show increased levels of arachidonic acid (20:4) in the CDAHFD and GAN livers, respectively, which are unaltered by MBOAT7 overexpression. (B,F) Total free fatty acid levels are largely unaltered in NASH livers compared to LF. (C-D, G-H) Abundance of hepatic acyl-CoA species and total measured acyl-CoA indicates that almost all acyl-CoA and particularly arachidonoyl-CoA (20:4) is decreased in the CDAHFD and GAN models of NASH, respectively, and not altered by MBOAT7 overexpression. Data presented as mean ± SEM, CDAHFD study: *n*=10 each; GAN study: LF n=10, GFP n=15, MBOAT7 n=18. Data analyzed by one-way ANOVA with Tukey’s post-hoc correction for multiple comparisons, \**p* < 0.05, \*\**p* < 0.001, \*\*\**p* < 0.001, \*\*\*\**p* < 0.0001.

The enzymes responsible for ligating long-chain fatty acids with CoA belong to the long-chain acyl-CoA synthetase (ACSL) family (Fig. 6A). Increased arachidonic acid levels and decreased arachidonoyl-CoA suggests decreased ACSL activity in NASH livers. There are several ACSL isoenzymes, with ACSL4 described as particularly important for arachidonic acid ligase activity (28). We first measured the hepatic gene expression of *Acsl1*, *Acsl3*, *Acsl4*, and *Acsl5* in these livers, which all displayed significantly reduced expression in livers from both NASH diets compared to LF except for *Acsl4* in the CDAHFD which was unchanged (Fig6B-I). Protein expression for ACSL1 and ACSL4 was significantly decreased in the CDAHFD model (Fig. 6J,L,M), while the GAN model of NASH decreased ACSL4 but not ACSL1 expression compared to LF (Fig. 6K,N,O). In summary, these data from two dietary NASH models suggests that arachidonoyl-CoA is decreased in NASH potentially due to decreased ACSL expression and activity.

**FIGURE 6.**
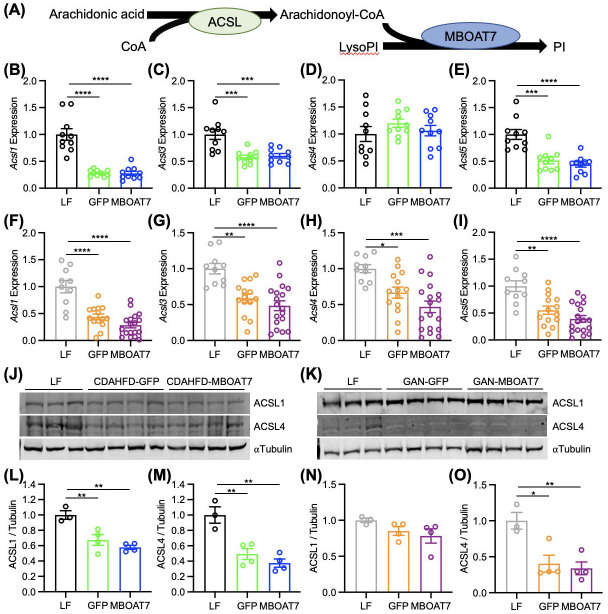
Decreased long-chain acyl-CoA synthetase (ACSL) expression limits arachidonoyl-CoA availability. (A) Schematic pathway of arachidonic acid incorporation into PI. (B-I) Hepatic gene expression of *Acsl1*, *Acsl3*, *Acsl4*, and *Acsl5* almost all significantly decreased in the CDAHFD and GAN models of NASH, and unaltered by MBOAT7 overexpression. (J,K) ACSL1 and ACSL4 expression is decreased in both the CDAHFD and GAN NASH models compared to LF. Data presented as mean ± SEM, CDAHFD study: *n*=10 each; GAN study: LF n=10, GFP n=15, MBOAT7 n=18. Data analyzed by one-way ANOVA with Tukey’s post-hoc correction for multiple comparisons, \**p* < 0.05, \*\**p* < 0.001, \*\*\**p* < 0.001, \*\*\*\**p* < 0.0001.

## Discussion

A common polymorphism near *MBOAT7* (rs641738), associates with increased hepatic steatosis and fibrosis (5–7), and is now widely considered a genetic risk allele for NASH (4, 29). This C>T variant decreases MBOAT7 expression (5), however another study suggests obesity decreases hepatic *MBOAT7* independent of the rs641738 polymorphism (10). In this present study, we mined publicly available genomic and lipidomic databases to confirm that livers from humans with obesity, NAFLD, or NASH display decreased MBOAT7 expression and activity compared to lean controls. A wealth of recent data from cell and rodent models suggests that MBOAT7-deficiency induces hepatic steatosis and NASH injury and fibrosis (10–14). The exact mechanism for increased steatosis and injury with MBOAT7 deficiency remains unclear, however several studies suggest increased SREBP1 activity and *de novo* lipogenesis (11, 14). If decreased MBOAT7 activity is a driving factor for NASH liver injury, we postulated that increasing MBOAT7 expression and activity would improve NASH. We addressed this question by overexpressing MBOAT7 in the liver with adeno-associated virus in two diet-induced models of NASH in mice. Unfortunately, most histologic measures of NASH were not improved by MBOAT7 in either model.

Since MBOAT7 is a lysophosphatidylinositol acyltransferase with a preference for arachidonic acid (8, 9), we performed lipidomic analyses to measure LPI and PI species in these livers to assess the functional implications of MBOAT7 overexpression. Both NASH models resulted in an increase in LPI and decreased PI suggestive of decreased MBOAT7 activity, despite no obvious decrease in MBOAT7 expression in these models. Two of the most abundant PI species contain arachidonic acid (36:4 and 38:4 PI), thus would be expected to be increased by MBOAT7, yet to our surprise, the levels of 36:4 and 38:4 PI were not rescued by MBOAT7. However, many other PI species, including non-arachidonate containing species, were significantly elevated by MBOAT7 overexpression. In agreement with these results, lipidomic analyses performed in MBOAT7 knockout models have identified non-arachidonate containing PI species that are reduced with MBOAT7 deficiency (10–12, 14). Thus, one conclusion that can be made from the present study is that *in vivo* MBOAT7 is not entirely specific to arachidonic acid.

In agreement with other studies (8, 10–14), we observed extremely few alterations in other hepatic phospholipids, cholesteryl esters, ceramides, or sphingomyelin species from modulating MBOAT7 expression. From these analyses we noted that arachidonate-containing (both 36:4 and 38:4) PE and PS were also significantly reduced by both NASH diets compared to LF control. This led us to hypothesize that NASH livers had limited availability of arachidonic acid to incorporate into phospholipids, potentially explaining why 36:4 and 38:4 PI were not enhanced by MBOAT7 overexpression. Indeed, although free arachidonic acid was significantly elevated, the activated form, arachidonoyl-CoA, which is a required intermediate for incorporation into phospholipids, was significantly reduced by both NASH diets compared to LF diet. To our knowledge hepatic arachidonoyl-CoA levels have not previously been measured in human or rodent models of NAFLD/NASH. Several rodent studies also identified increased free arachidonic acid in NAFLD/NASH vs control livers (30, 31), however human data suggests decreased free arachidonate in NASH (30, 32–34). Nevertheless, reduced arachidonate content in hepatic phospholipids and/or triglycerides has been observed in several studies of human NASH, in agreement with our current findings (20, 21, 30, 34). Increased omega oxidation products of arachidonate have been demonstrated in plasma lipidomic studies of NASH patients, suggesting increased disposal may reduce arachidonate availability for incorporation into phospholipid species (35). ACSL4 is believed to play a specific role in arachidonoyl-CoA synthesis (28). Hepatic ACSL4 was significantly decreased in both of our NASH models and is also decreased by high-fat diet feeding in mice (36). Interestingly, free arachidonic acid causes ACSL4 to be ubiquitinated and degraded, leading to decreased ACSL4 protein expression (36, 37). However, this decrease in hepatic ACSL4 in NAFLD/NASH may be restricted to rodent models, as increased ACSL4 has been identified in human NAFLD (38, 39). And although we and others (36) observe decreased ACSL4 in rodent models of NAFLD/NASH, a recent report described that ACSL4 deletion or pharmacologic inhibition in mice protected from NASH primarily by enhancing fat oxidation (39). Thus, more studies are required to gain a better understanding of the importance of ACSL4 and arachidonoyl-CoA in both humans and rodent models of NASH.

In summary, in this study we observe lipid changes suggesting a loss of MBOAT7 activity in two diet-induced models of murine NASH, but found hepatic MBOAT7 overexpression was unable to substantially improve NASH pathology although markers of liver injury were improved. This lack of measurable histologic improvement may be due to the inability of MBOAT7 to enhance the major arachidonoylated PI (38:4 and 36:4) species, likely due to the low arachidonoyl-CoA levels from decreased ACSL4 expression in NASH livers. These findings suggest that MBOAT7 may not be an actionable target in NASH but set the stage for further studies related to the activity of ACSL4 and arachidonoyl-CoA in NASH.

## List of abbreviations

AAV8: adeno-associated virus serotype 8
ACSL: long-chain acyl-CoA synthetase
ALT: alanine aminotransferase
AST: aspartate aminotransferase
CDAHFD: choline-deficient amino acid defined high-fat diet
GAN: Gubra Amylin NASH diet
GEO: gene expression omnibus
GFP: green fluorescent protein
GTT: glucose tolerance test
HO: healthy obese
LDHA: lactate dehydrogenase A
LF: low-fat diet
LPC: lysophosphatidylcholine
LPI: lysophosphatidylinositol
MBOAT7: membrane-bound O-acyltransferase 7
NAS: NAFLD activity score
NC: normal control
NEFA: non-esterified fatty acid
PC: phosphatidylcholine
PE: phosphatidylethanolamine
PI: phosphatidylinositol
PS: phosphatidylserine
TAG: triglyceride
TBG: thyroid binding globulin
QC: quality control
SREBP1: sterol regulatory element binding protein 1

## Acknowledgements

We thank Drs. Hiroyuki Arai and Nozomu Kono for the gracious gift of the MBOAT7 antibody.

## Author Contributions

M.C.S.: data curation, formal analysis, investigation, methodology, writing – review and editing; K.D.P.: data curation, formal analysis, investigation, methodology, writing – review and editing; T.H.: data curation, formal analysis, investigation, methodology, writing – review and editing; D.R.K.: data curation, formal analysis, investigation, methodology, writing – review and editing; M.P.: data curation, formal analysis, investigation, methodology, writing – review and editing; B.F.: formal analysis, software, writing – review and editing; C.J.A.: data curation, formal analysis, investigation, methodology, supervision, validation, writing – review and editing; D.H.C.: formal analysis, writing – review and editing; B.U.: conceptualization, data curation, formal analysis, investigation, methodology, project administration, supervision, visualization, writing – review and editing; D.A.F.: data curation, formal analysis, methodology, resources, project administration, supervision, writing – review and editing; B.A.N.T.: conceptualization, project administration, supervision, visualization, writing – review and editing; K.S.M.: conceptualization, data curation, formal analysis, funding acquisition, investigation, methodology, project administration, supervision, visualization, writing – original draft, writing – review and editing.

