## Supplemental Table 1 for "Enhancing Hepatic MBOAT7 Expression Does Not Improve Nonalcoholic Steatohepatitis in Mice"

**Supplemental Table 1. List of oligonucleotide primers for qPCR.**

| Gene name | FWD primer (5' - 3') | REV primer (5' - 3') |
|---|---|---|
| *Acsl1* | TGGGGTGGAAATCATCAGCC | CACAGCATTACACACTGTACAACGG |
| *Acsl3* | GGGACTACAATACCGGCAGAGT | AATAGCCACCTTCCTCCCAGTT |
| *Acsl4* | AAATGCAGCCAAATGGAAAG | CACTCTGCAGTTCACTTCAA |
| *Acsl5* | ATCTGCCTCCTGACGTTTGG | GCTCCTCCCTCAATCCCCAC |
| *Acta2* | GTCCCAGACATCAGGGAGTAA | TCGGATACTTCAGCGTCAGGA |
| *Col1a1* | GCTCCTCTTAGGGGCCACT | CCACGTCTCACCATTGGGG |
| *Col3a1* | CTGTAACATGGAAACTGGGGAAA | CCATAGCTGAACTGAAAACCACC |
| *Mboat7* | ATCCGTAACATCGACTGCTATGG | CGGAAAGGTGCGCTCTTGTA |
| *Mmp2* | ACCTGAACACTTTCTATGGCTG | CTTCCGCATGGTCTCGATG |
| *Timp1* | CGAGACCACCTTATACCAGCG | ATGACTGGGGTGTAGGCGTA |
| *Rplp0* | AGATTCGGGATATGCTGTTGGC | TCGGGTCCTAGACCAGTGTTC |
