## Supplemental Table 2 for "Enhancing Hepatic MBOAT7 Expression Does Not Improve Nonalcoholic Steatohepatitis in Mice"

**Supplemental Table 2. List of antibodies used for western blot analyses.**

| Antibody | Host Species | Source | Catalog/Ref. # |
|---|---|---|---|
| $\alpha$-Tubulin | Rabbit | Cell Signaling Technology | 2125S |
| ACSL1 | Rabbit | Proteintech | 13989-1-AP |
| ACSL4 | Rabbit | MilliporeSigma | SAB2100035 |
| Calnexin | Rabbit | Enzo | ADI-SPA-860 |
| LDHA | Rabbit | Cell Signaling Technology | 2012S |
| MBOAT7 | Rat | Custom made monoclonal (FT10) | PMID: 23097495 |
| Anti-Rabbit 680RD | Donkey | LiCor | 926-69073 |
| Anti-Rabbit 800CW | Donkey | LiCor | 926-32213 |
| Anti-Rat 800CW | Goat | LiCor | 926-32219 |
