## Supplementary material for "Enhancing Hepatic MBOAT7 Expression Does Not Improve Nonalcoholic Steatohepatitis in Mice": Fig. S1

Supplemental Figure 1

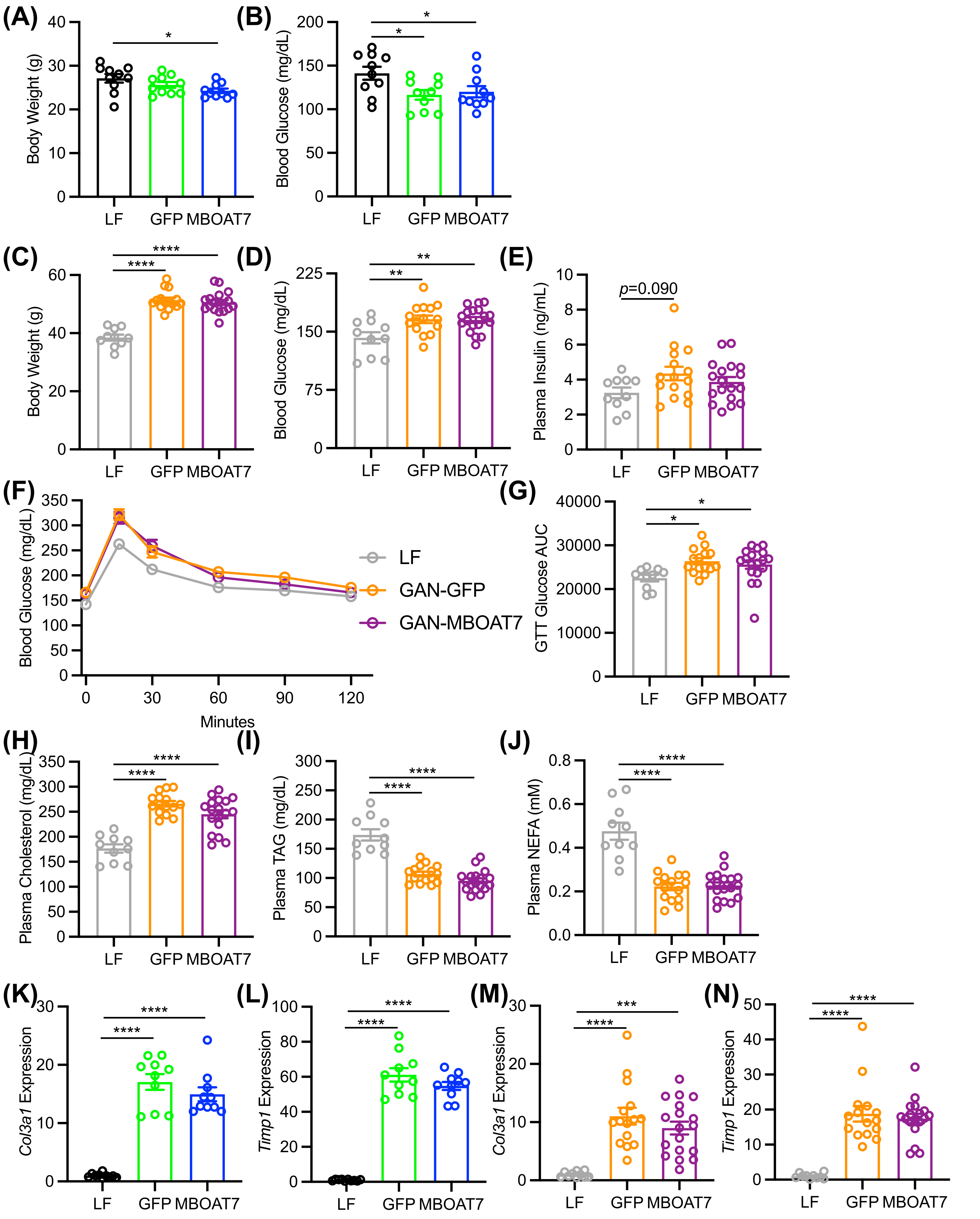

**Figure S1** Body weights, plasma glycemia and lipids, and hepatic gene expression data in CDAHFD and GAN models of NASH. (A) Body weights in LF vs CDAHFD-fed mice. (B) Blood glucose concentrations decreased in CDAHFD-fed compared to LF-fed mice, and not altered by MBOAT7. (C) Body weight increased by GAN diet compared to LF, not altered by MBOAT7 overexpression. (D) Blood glucose concentrations increased in GAN-fed compared to LF-fed mice, and not altered by MBOAT7. (E) Plasma insulin concentrations not altered by GAN compared to LF diet. (F and G) Blood glucose excursions during an i.p. glucose tolerance test and glucose area under the curve increased by GAN diet compared to LF, but not altered by MBOAT7 overexpression. (H-J) Plasma cholesterol, TAG, and NEFA in the GAN-model of NASH compared to LF diet. (K and L) *Col3a1* and *Timp1* expression are increased in CDAHFD livers compared to LF, but not improved by MBOAT7 overexpression. (M and N) *Col3a1* and *Timp1* expression are increased in GAN livers compared to LF, but not improved by MBOAT7 overexpression. Data presented as mean ± SEM, CDAHFD study: *n*=10 each; GAN study: LF n=10, GFP n=15, MBOAT7 n=18. Data analyzed by one-way ANOVA with Tukey's post-hoc correction for multiple comparisons, *$p < 0.05$, **$p < 0.001$, ***$p < 0.001$, ****$p < 0.0001$.
