## Supplementary material for "Enhancing Hepatic MBOAT7 Expression Does Not Improve Nonalcoholic Steatohepatitis in Mice": Fig. S2

Supplemental Figure 2

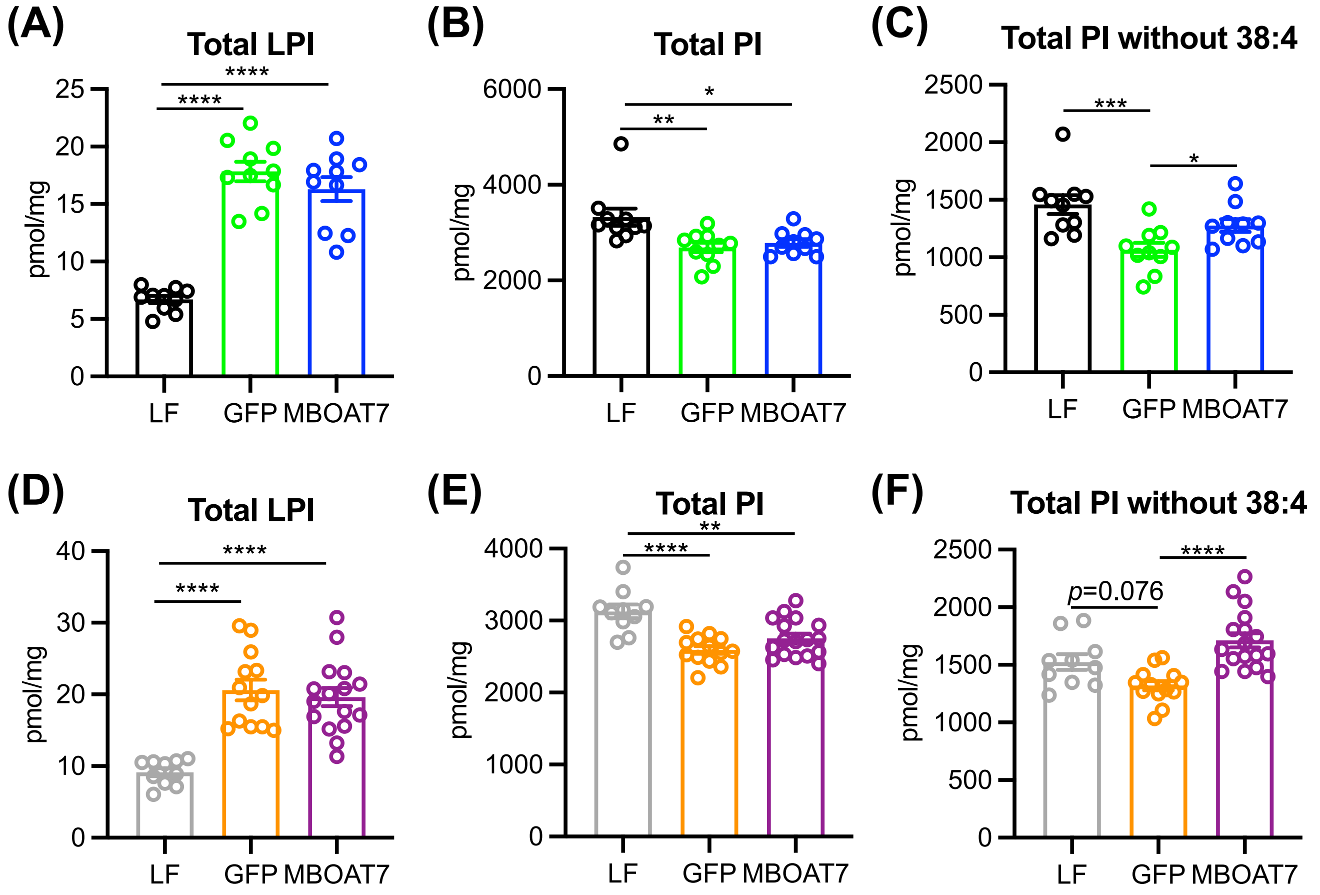


**Figure S2** Total hepatic lysophosphatidylinositol (LPI) and phosphatidylinositol (PI) in CDAHFD and GAN models of NASH. (A) Total LPI concentrations are increased in CDAHFD compared to LF livers, and not altered by MBOAT7 overexpression. (B) Total PI concentrations are decreased in CDAHFD compared to LF livers, and not altered by MBOAT7 overexpression. (C) If the abundant 38:4 PI is excluded, total PI concentrations are decreased in CDAHFD GFP compared to LF livers, and significantly increased by MBOAT7 overexpression. (D) Total LPI concentrations are increased in GAN compared to LF livers, and not altered by MBOAT7 overexpression. (E) Total PI concentrations are decreased in GAN compared to LF livers, and not altered by MBOAT7 overexpression. (F) If the abundant 38:4 PI is excluded, total PI concentrations are decreased in GAN GFP compared to LF livers, and significantly increased by MBOAT7 overexpression. Data presented as mean ± SEM, CDAHFD study: $n$=10 each; GAN study: LF n=10, GFP n=15, MBOAT7 n=18. Data analyzed by one-way ANOVA with Tukey's post-hoc correction for multiple comparisons, *$p < 0.05$, **$p < 0.001$, ***$p < 0.001$, ****$p < 0.0001$.
