## Supplementary material for "Enhancing Hepatic MBOAT7 Expression Does Not Improve Nonalcoholic Steatohepatitis in Mice": Fig. S3

Supplemental Figure 3

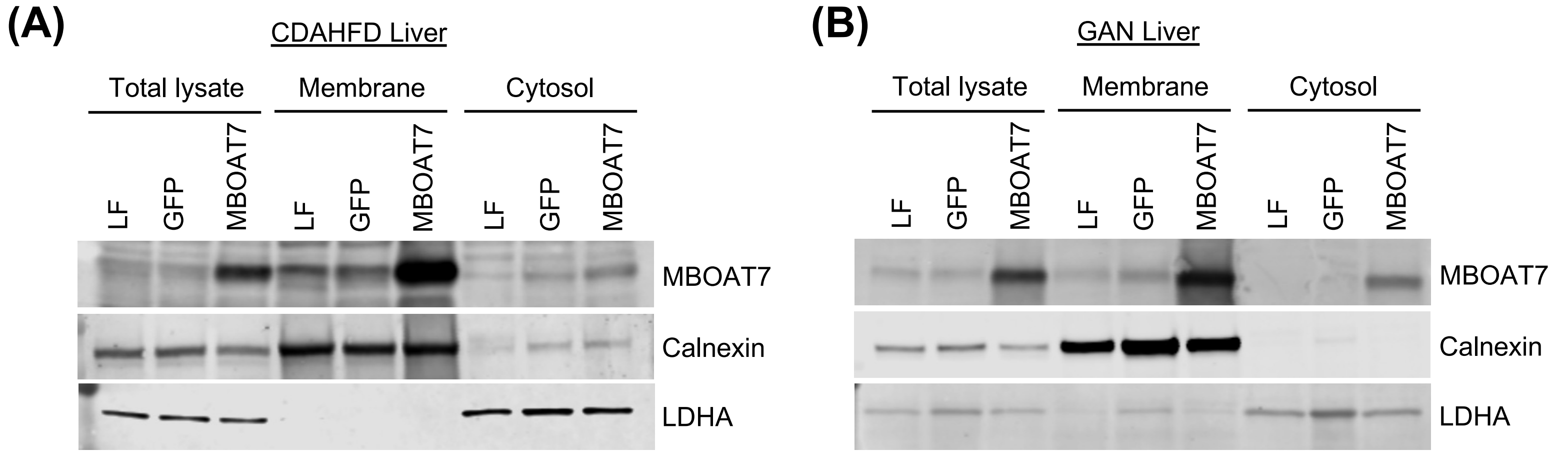


**Figure S3** MBOAT7 localization in membrane vs cytosolic liver fractions. (A and B) Western blots from liver tissue from CDAHFD and GAN studies, respectively, homogenized into total lysate or purified into membrane and cytosolic fractions by differential centrifugation. MBOAT7 is predominantly localized to the membrane fraction, marked by the endoplasmic reticulum marker calnexin. A small amount of the overexpressed MBOAT7 appears in the cytosolic fraction, marked by lactate dehydrogenase A expression.
