## Supplementary material for "Enhancing Hepatic MBOAT7 Expression Does Not Improve Nonalcoholic Steatohepatitis in Mice": Fig. S4

Supplemental Figure 4

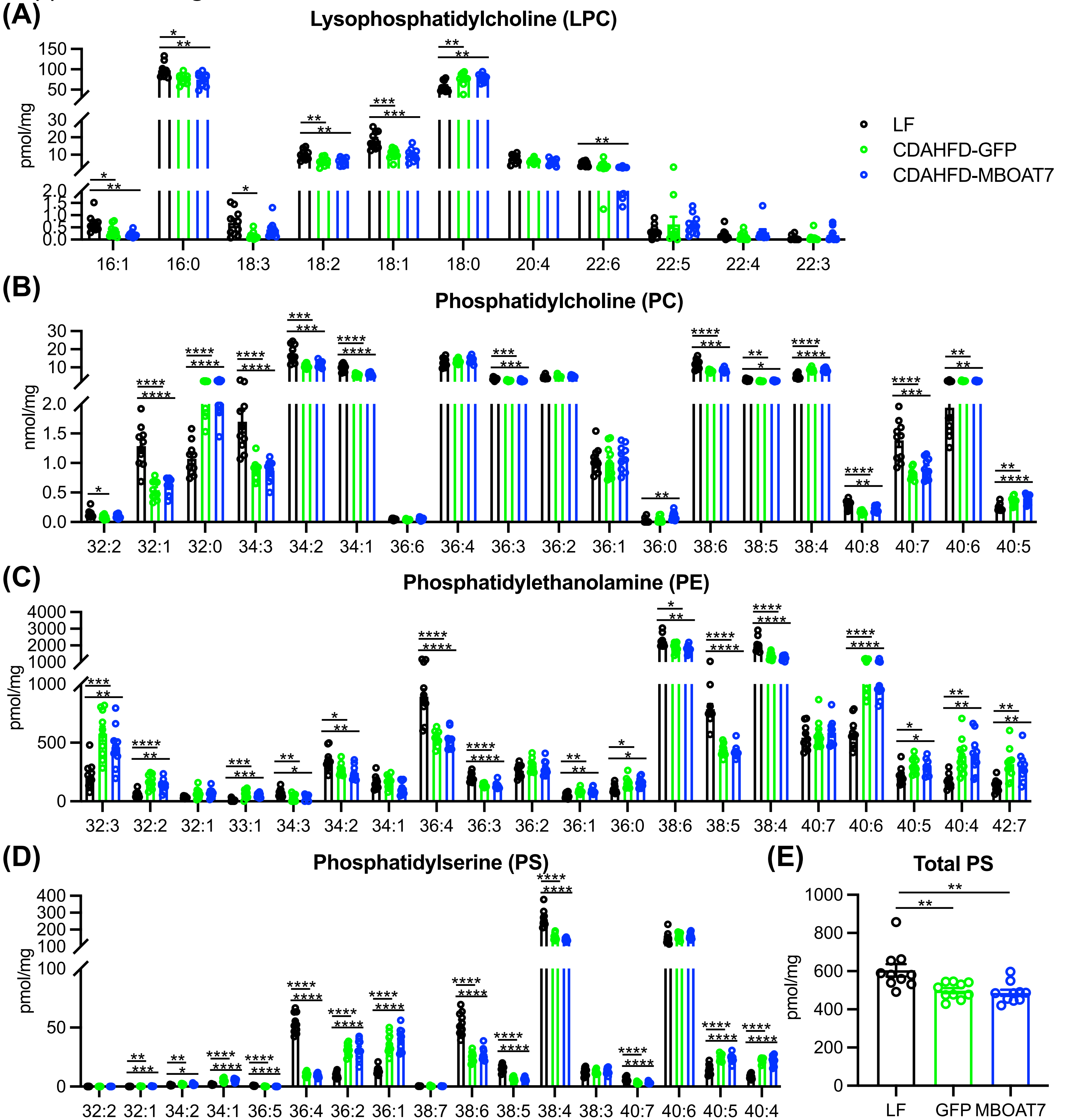


**Figure S4** Hepatic lysophosphatidylcholine (LPC), phosphatidylcholine (PC), phosphatidylethanolamine (PE), and phosphatidylserine (PS) in the CDAHFD model of NASH. (A-D) Hepatic LPC, PC, PE, and PS molecular species concentrations in CDAHFD compared to LF livers, with no significant changes between MBOAT7 overexpression and GFP control. (E) Total PS levels are decreased by CDAHFD compared to LF, and unaltered by MBOAT7 overexpression. Data presented as mean ± SEM, *n*=10 each. Data analyzed by one-way ANOVA with Tukey's post-hoc correction for multiple comparisons, **p* < 0.05, ***p* < 0.001, ****p* < 0.001, *****p* < 0.0001.
