## Supplementary material for "Enhancing Hepatic MBOAT7 Expression Does Not Improve Nonalcoholic Steatohepatitis in Mice": Fig. S5

Supplemental Figure 5

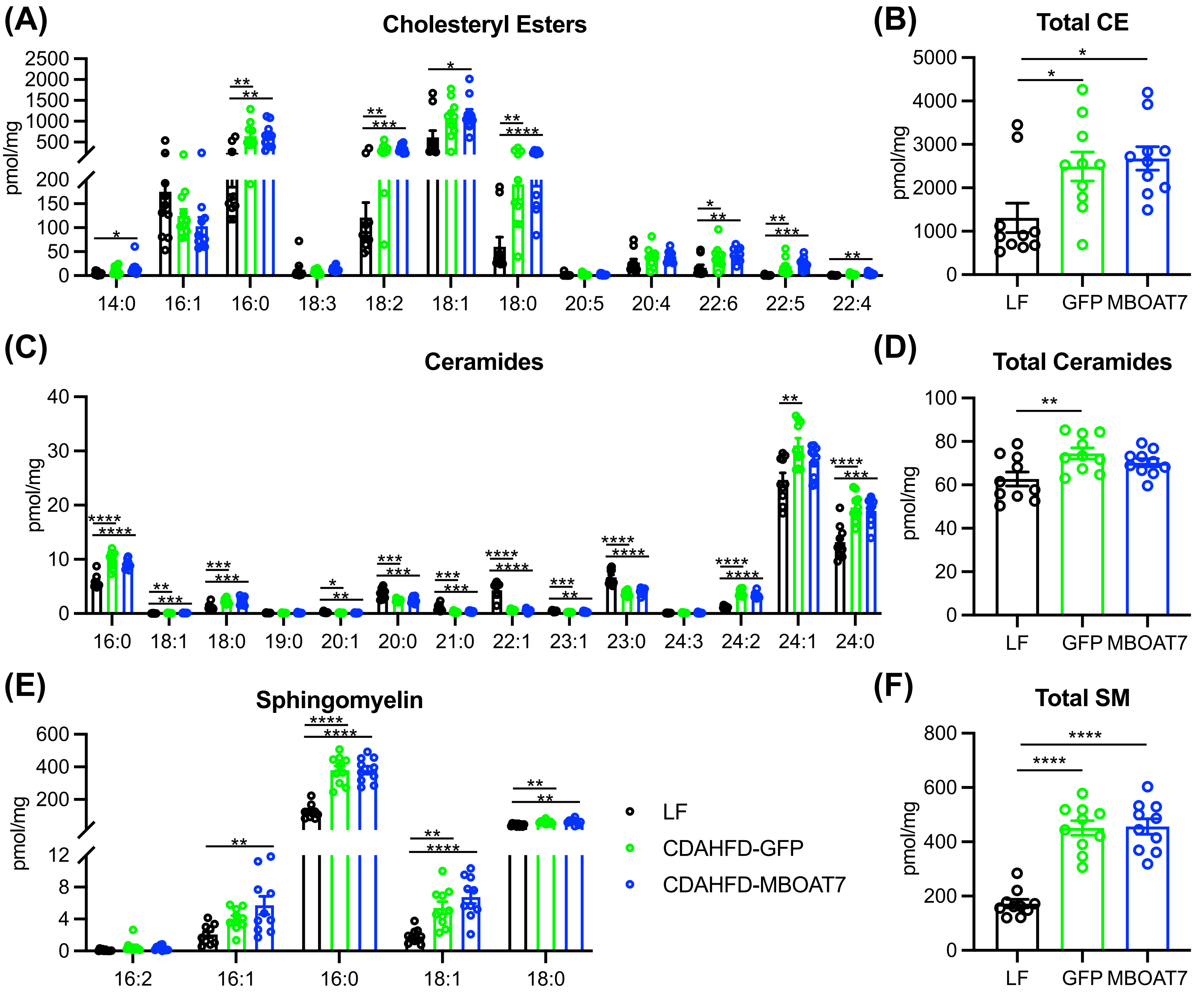


**Figure S5** Hepatic cholesteryl ester (CE), ceramide, and sphingomyelin (SM) concentrations in the CDAHFD model of NASH. (A and B) Individual CE species and total hepatic CE in CDAHFD compared to LF diet livers, with no significant alterations from MBOAT7 overexpression. (C and D) Individual ceramide species and total hepatic ceramides in CDAHFD compared to LF diet livers, with no significant alterations from MBOAT7 overexpression. (E and F) Individual SM species and total hepatic SM in CDAHFD compared to LF diet livers, with no significant alterations from MBOAT7 overexpression. Data presented as mean ± SEM, $n$=10 each. Data analyzed by one-way ANOVA with Tukey's post-hoc correction for multiple comparisons, *$p < 0.05$, **$p < 0.001$, ***$p < 0.001$, ****$p < 0.0001$.
