## Supplementary material for "Enhancing Hepatic MBOAT7 Expression Does Not Improve Nonalcoholic Steatohepatitis in Mice": Fig. S6

Supplemental Figure 6

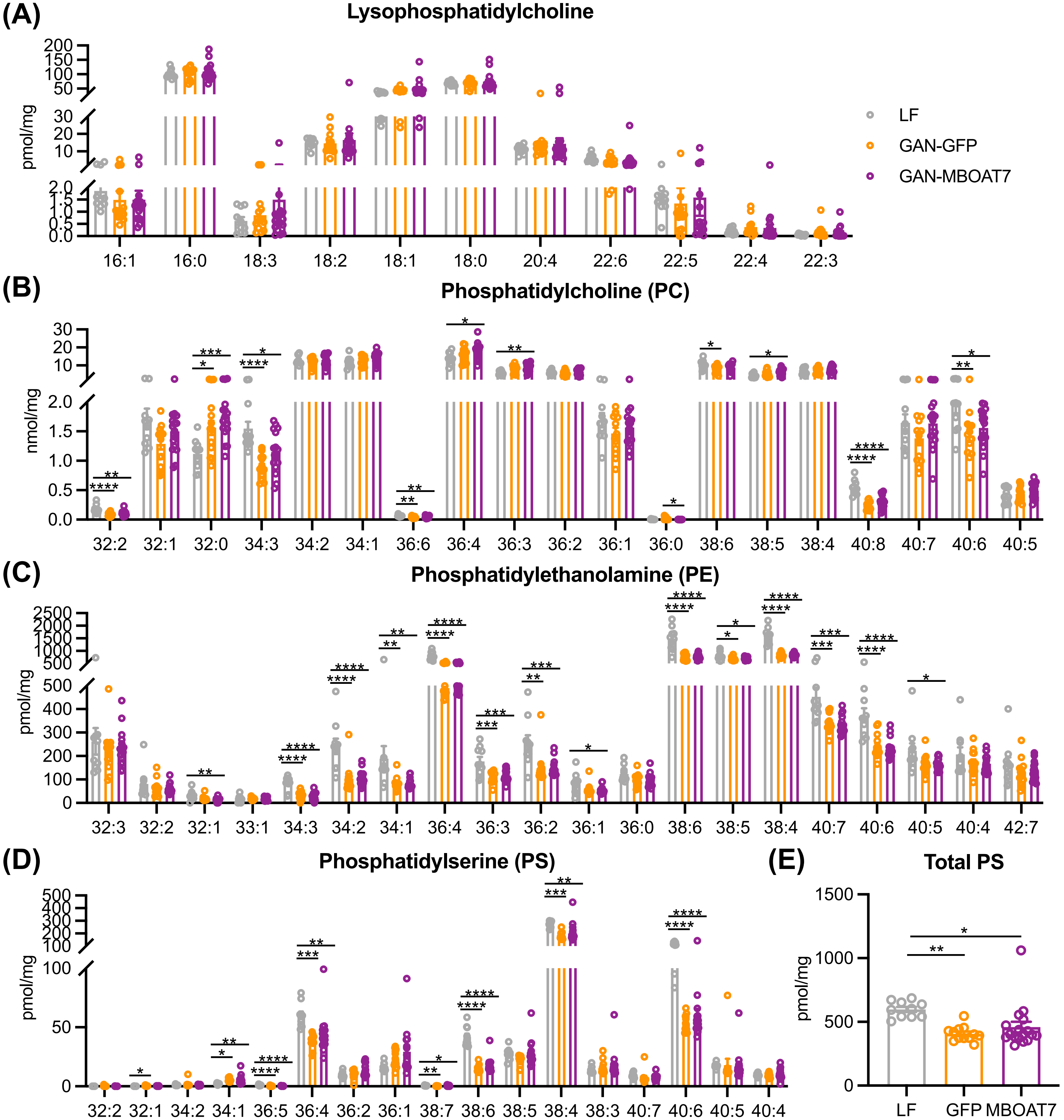


**Figure S6** Hepatic lysophosphatidylcholine (LPC), phosphatidylcholine (PC), phosphatidylethanolamine (PE), and phosphatidylserine (PS) in the GAN model of NASH. (A-D) Hepatic LPC, PC, PE, and PS molecular species concentrations in GAN compared to LF livers, with very few significant changes between MBOAT7 overexpression and GFP control. (E) Total PS levels are decreased by GAN compared to LF, and unaltered by MBOAT7 overexpression. Data presented as mean ± SEM, LF n=10, GFP n=15, MBOAT7 n=18. Data analyzed by one-way ANOVA with Tukey's post-hoc correction for multiple comparisons, *$p < 0.05$, **$p < 0.001$, ***$p < 0.001$, ****$p < 0.0001$.
