## Supplementary material for "Enhancing Hepatic MBOAT7 Expression Does Not Improve Nonalcoholic Steatohepatitis in Mice": Fig. S7

Supplemental Figure 7

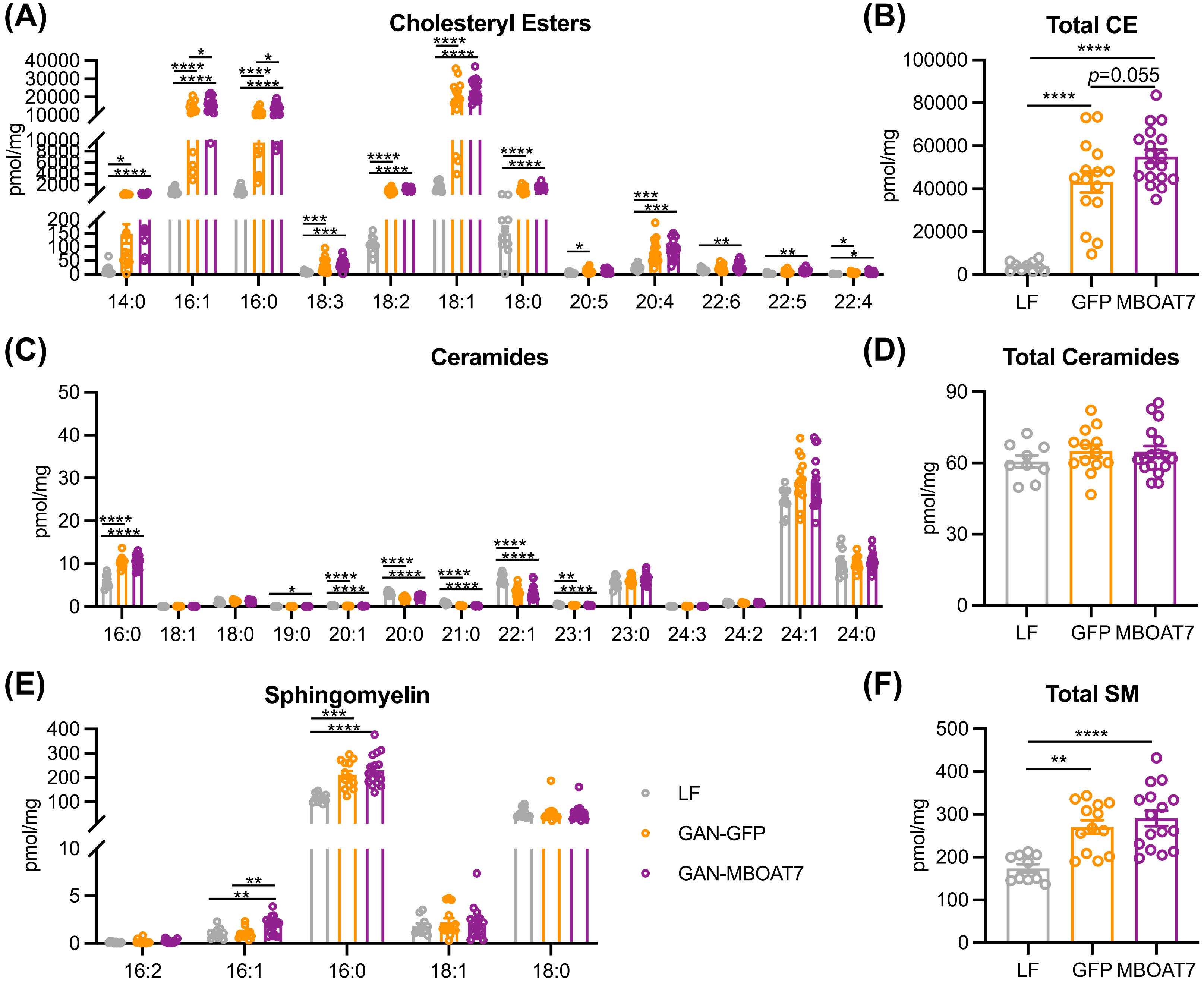


**Figure S7** Hepatic cholesteryl ester (CE), ceramide, and sphingomyelin (SM) concentrations in the GAN model of NASH. (A and B) Individual CE species and total hepatic CE in GAN compared to LF diet livers. (C and D) Individual ceramide species and total hepatic ceramides in GAN compared to LF diet livers, with no significant alterations from MBOAT7 overexpression. (E and F) Individual SM species and total hepatic SM in GAN compared to LF diet livers. Data presented as mean ± SEM, LF n=10, GFP n=15, MBOAT7 n=18. Data analyzed by one-way ANOVA with Tukey's post-hoc correction for multiple comparisons, *$p < 0.05$, **$p < 0.001$, ***$p < 0.001$, ****$p < 0.0001$.
